# Contribution of Notch/Wnt signaling modulation in reactive astrocyte reparative response after brain injury

**DOI:** 10.1101/2022.12.20.521209

**Authors:** Lina Maria Delgado-Garcia, Julia C. Benincasa, Natália Rodrigues Courbassier, Tais Novaki Ribeiro, Marimélia Porcionatto

## Abstract

After a Traumatic Brain Injury (TBI), the neural network activates a reparative response seeking to restore homeostasis. Astrocyte reactivation is an essential component of this response. The injury creates a temporal microenvironment where neurogenic signaling molecules regulate cell fate decisions of neocortical neural progenitors. Likewise, astrocyte reactivation triggers a transcriptional-proliferative program where neurogenic signaling molecules play crucial roles. However, precise molecular mechanisms are context-specific and are not fully understood. Here we studied cellular and molecular aspects of reactive astrocytes response after Notch-Wnt neurogenic signaling modulation. Our results provide new evidence of cortical Notch-Wnt signaling activation after TBI. Reactive astrocytes in the core of Notch signaling showed a differential aggregated distribution. *In vitro*, Notch inhibition promoted a neural precursor profile and might increase the number of cells committed in a proliferative response. Finally, we found an indirect co-regulation of Wnt-Shh signaling in BHLH-Notch target genes and a Notch-supportive effect in Wnt-Shh signaling activation.

## Introduction

After any neurological condition, such as a Traumatic Brain Injury (TBI), the neural network activates a reparative response to restore homeostasis and minimize tissue damage. Rapidly, three overlapping stages are recognized, from cell death and inflammation to cell proliferation and tissue remodeling (Burda et Sofroniew., 2014; Wahane et Sofroniew., 2021). Annually, 69 million people suffer from minor brain damage such as a concussion to moderate and severe conditions such as contusions and penetrating injuries (Dewan et al., 2018). For the last two cases, TBI often results in life-changing conditions. TBI might impact higher motor and cognitive functions, such as memory, movement, and sensation. Other possible complications include long-term or prolonged motor and cognitive deficits and the development of psychiatric, neurodegenerative, and neuroinflammatory disorders (Bramlett et Dietrich., 2015).

At the physiopathological level, astrocyte reactivation is an essential component in brain reparative response to a TBI (Pekny et Pekna, 2014; Verkhratsky, et Parpura, 2016; Verkhratsky et Nedergaard, 2018). Mediators of astrocyte reactivity like ATP, purines and pyrimidines are released by dying or injured cells (Abbracchio et Ceruti, 2006) around the lesion core, giving rise to an astrocytés paracrine activation loop (Buffo et al., 2010). It’s known that these reactive astrocytes are characterized by upregulation of cytoskeletal proteins such as glial fibrillary acidic protein, GFAP, neuroepithelial stem cell protein, Nestin and Vimentin, and the hypertrophy of the soma and processes. Subpopulations of cortical reactive astrocytes at juxtavascular sites proliferate while others reactive astrocytes populations elongate their processes towards the lesion site (Bardehle et al., 2013; Liddlelow et Barres, 2017; Escartin et al., 2021). Although reactive astrocytes aim at protecting tissue homeostasis, some studies have shown detrimental effects associated with astrogliosis. Extracellular matrix components such as chondroitin (CSPG) and keratan sulfate proteoglycans produced by reactive astrocytes at the lesion border inhibit axonal growth (Silver et Miller, 2004) and impairs neural stem migration to the injury site (Galindo et al., 2018). However, glial scar ablation is insufficient to promote axonal re-generation (Anderson et al., 2016), and deficient GFAP−/− Vim−/− mice showed poor synaptic restoration after entorhinal cortex lesion (Wilhelmsson et al., 2004). Thus, astrocyte reactivation response to a damage depends on injurýs severity, extension, anatomical location, and mechanical properties (Zamanian et al., 2012; Burda et al., 2016; Adams et Gallo, 2018). Indeed, glial scar appears only after a strong stimulus. In that case, astrocytes acquire a reactive-proliferative program that creates a non-reversible glial scar to restore homeostasis efficiently and hinder inflammation to adjacent healthy tissue (Anderson et al., 2016). On the other hand, a mild, non-penetrating stimulus promotes a temporal astrocyte reactivation program, non-proliferative and reversible (Burda et al., 2016). Certainly, there is still a vast field of research to understand astrocyte reactivation from the molecular to the physiological level and how it can be beneficial or detrimental to brain recovery.

In the adult brain, neurogenic signaling molecules modulate multipotent neural precursors in restricted areas called neurogenic niches to replace lost neurons (Urbán et Guillemont, 2014; Choe et al., 2015; Bonfanti, 2016). These morphogens and signaling molecules create concentration gradients while it spreads through the tissue. Thus, signaling molecules play a role in a concentration- and time-dependent manner promoting gene expression and activation of neurogenic programs (autorenewal, proliferation, and differentiation). Moderate changes are sufficient to significantly shift the celĺs response to the same molecule (Ashe et Briscoe, 2006). In the context of brain damage, the injury creates a temporal microenvironment for cell proliferation and tissue remodeling, resembling the neurogenic niches. Several soluble and membrane factors have been reported, including neurogenic signaling molecules growth factors, and neurotrophins (Roll et Faissner, 2014; Burda et Sofroniew, 2014). Likewise, reactive astrocytes overexpress growth factors and their receptors to provide trophic support for damaged neurons, oligodendrocytes, and progenitor cells (Planas et. al, 1998).

Notch, Wnt, and Shh are neurogenic signaling molecules widely used by neocortical neural progenitors during brain development. In the adult brain, these molecules persist mainly in the neurogenic niches. Notch is a transmembrane protein that signals through cell-cell interactions. Briefly, when the Notch receptor binds to any of their ligands, Jagged or Delta, the γ-secretase enzyme cleaves the cytoplasmic portion of Notch and releases the Notch intracellular domain NICD. That is because ligand binding to Notch exerts a pulling force, which exposes the Notch cleavage site for ADAM10 or ADAM17. Then, NICD is translocated to the nucleus, binds the transcription factor RBPJK and recruits the coactivator Mastermind-like MAML creating the complex NICD/RBPJK/MAML that promotes the expression of BHLH transcriptional repressors HES (HES 1, HES 3, HES 5) and HEY. In progenitor cells, HES factors repress the expression of target proneural genes such as MASH1 (ASCL1) and NEUROGENIN2 (Imayoshi et al., 2010, 2013; Urbán et Guillemont, 2014; Choe et al., 2015; Kageyama et al., 2015). On the other hand, Wnt and Shh neurogenic signaling pathways are soluble factors, that form extracellular gradients in the neurogenic niches. Wnt signaling comprises a wide variety of receptors, ligands, inhibitors, and target genes. In mice, there are at least ten types of frizzled receptors and several coreceptors. Wnt signaling has a canonical (dependent of ß-catenin) and a non-canonical (ß-catenin independent) activation pathway. In the absence of Wnt signaling, ß-catenin integrates into the proteasomal degradation complex axin/APC/GSK3β. Thus, in the canonical pathway, the Wnt ligand binds the receptor frizzled, which favors the stabilization and accumulation of ß-catenin in the cytoplasmic compartment. Activated ß-catenin is translocated to the nucleus and binds the transcription factor complex T-cell factor/lymphoid enhancer-binding factor, TCF/LEF to regulate a wide variety of target genes, including BHLH and homeobox proneural genes (NEUROGENIN2, NEUROD1 and PROX1, Gordon et Nusse, 2006; Angers et Moon 2009; Kuwabara et al., 2009; Karalay et al., 2011; Varela-Nallar et Inestrosa, 2013; Choe et al., 2015). In the case of Shh signaling, binding of Shh ligands to the receptor patched, Ptch1; abolish the effect of Ptch1 on the G-protein receptor Smooth-ened, Smo. Then, Smo activation stabilizes cytoplasmic GLI (GLI1, GLI2, GLI3) and exposes the GLI carboxy-terminal domain, blocking their degradation in the proteasomal complex PKA/GSK3β/CKI and ubiquitin-ligase E3, and promoting their translocation to the nucleus. GLI proteins act as a transcription factor regulating homeobox and BHLH genes as OLIG2 (Palma et al., 2004, 2005; Pan et al., 2006; Wang et al., 2007; Urbán et Guillemont., 2014; Choe et al., 2015). In addition to the cited BHLH and homebox genes, Wnt and Shh signaling are well-known regulators of cell cycle regulators such as CCND1, CCND2, and CCNDE (Coyle-Rink et al., 2002; Ruiz I Altaba et al., 2002; Oliver et al., 2003; Komada et al., 2013; Araújo et al., 2014; Liu et al., 2018). Collectively, at the cellular level, extracellular signaling molecules bind to their corresponding receptor in the membrane of the cell which activates a cascade of intracellular protein complexes, producing spatio-temporal changes in gene expression and amplification of the signal.

The combinatorial effect of Notch, Wnt, and Shh neurogenic signaling pathways modulate crucial processes such as progenitor expansion, proliferation, migration, and neural differentiation (Urbán et Guillemont., 2014; Choe et al., 2015). Astrocyte reactivation triggers a transcriptional proliferative program where neurogenic signaling molecules play crucial roles, however, precise molecular mechanisms are context-specific and are not fully understood. Here we studied cellular and molecular aspects of reactive astrocytes response after Notch-Wnt neurogenic signaling modulation. We used an in *vivo*/*in vitro* model of brain injury and astrocyte reactivation. Our results provide new evidence of cortical Notch (NICD) and Wnt (active β-catenin) signaling activation after TBI. Reactive astrocytes in the core of Notch signaling presented a differential aggregated distribution. *In vitro*, Notch inhibition promoted a neural precursor profile (proliferation and self-renewal) and might increase the number of cells committed in a proliferative response. Finally, we found an indirect co-regulation of Wnt-Shh signaling in BHLH-Notch target genes and a Notch-supportive effect in Wnt-Shh signaling activation. Dissecting the role of supportive molecules in brain damage is crucial for our current comprehension of the mechanisms leading to brain repair and regeneration and the development of novel therapeutic interventions.

## Results

### TBI triggered neural repair mechanisms and specific signaling activation of neurogenic pathways

Adult C57bl/6j mice were submitted to a model of brain damage by TBI in the somatosensorial cortex. Three days post-injury (3dpi), we performed immunohistochemical analysis of reactive astrocyte response, proliferation and neurogenic signaling activation. Immunohistochemical markers of neuroinflammation and reactive astrocytes response included glial fibrillary acid protein (GFAP), S100 calcium-binding protein β (S100β) and, Galectin-3 (Gal3). GFAP and S100β are the most frequent markers and hallmark proteins in astrocyte reactivation (Verkhratsky et al., 2019; Escartin et al., 2021). Gal3 is a protein widely present in the central nervous system (CNS) that increases during neuroinflammation and diverse pathological conditions (Nio-Kobayashi., 2017; Yoo et al., 2017; Rahimian et al., 2018; Ribeiro et al., 2021). We used proliferation/cell cycle markers Ki67 and BRDU to identify newly proliferated cells and potential areas of neural cell genesis. Ki67 is a nuclear protein present during all cell cycle stages (Scholzen et al., 2000). On the other hand, Bromodeoxyuridine (BrdU) is a thymidine analog incorporated in the DNA of dividing cells during the S-phase of the cell cycle (Taupin, 2007).

We identified increased reactive astrocyte response in the ipsilateral-somatosensorial cortex. Reactive astrocytes showed enlarged cell bodies, hypertrophic primary and secondary branches, and immunolabeling of GFAP and S100β proteins. Quantitative analysis showed GFAP upregulation (mean gray value). Gal3 immunolabeling was preferentially distributed in the ipsilateral cortex, where half of the total population of reactive astrocytes exhibited GFAP/Gal3 overlapping (Figure 1A). There was progressive increase in newly proliferated Ki67+ cells from day 1 to day 3 (1–3-day post injury, dpi). Cells expressing Ki67+ and BRDU+ markers were found almost exclusively in the ipsilateral region. Ki67+ newly-proliferative cells represented the tenth part of the total population. Moreover, half of the Ki67+ total population of newly proliferated cells corresponded to GFAP reactive astrocytes (%Ki67/GFAP), which in turn, represented a third part of the total of reactive astrocytes population (Figure 1B).

**Figure 1.**
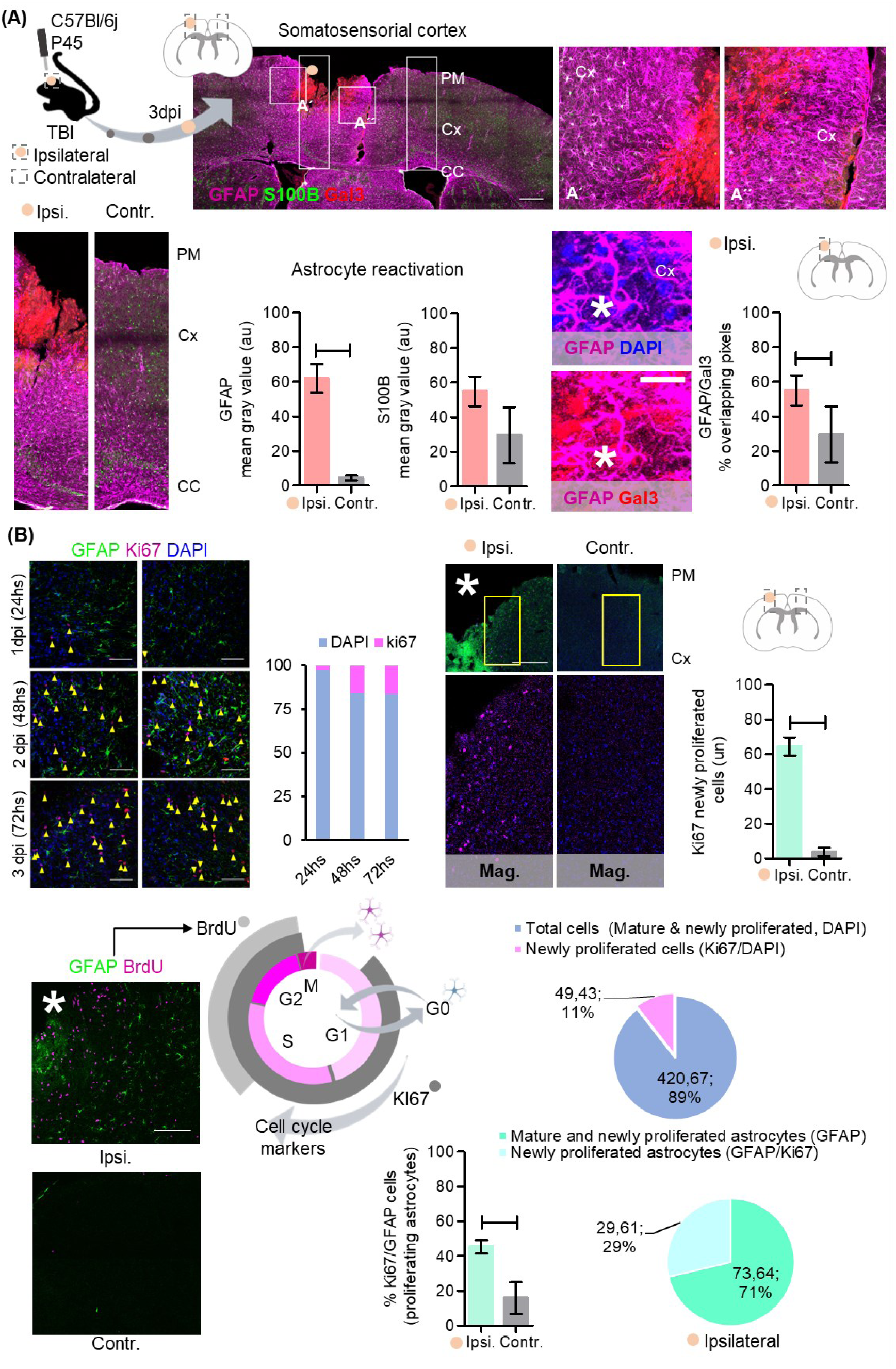
Astrocyte reactivation response and proliferation. (A). Experimental design and analysis. Adult (P45) C57Bl/6j mice were submitted to a model of brain damage by traumatic brain injury (TBI) in the somatosensorial cortex to explore astrocyte reactivation, proliferation, and neurogenic signaling activation. Representative images of reactive astrocytes and GFAP, S100β, and Gal3 immunolabeling. Quantitative analysis of GFAP and S100β immunolabeling and Gal3 protein distribution in reactive astrocytes (GFAP/Gal3 overlapping pixels). *Representative image of GFAP/Gal3 reactive astrocyte in the ipsilateral cortex. (B). Representative images of the proliferative response 1-, 2- and 3-days post-injury (dpi). Newly proliferated cells (Ki67+ and BrDU+) were abundant in the ipsilateral cortex. We found an increased proportion in the total number of cells (Dapi) and the number of newly-proliferated cells (Ki67+). Newly-proliferated astrocytes (GFAP/Ki67+) were about a third of the total. N=3 biological replicates. Values with p < 0.05 were considered statistically significant. Scale bar 100 μm and 20 μm* in A, 50 μm in B, up and down, left, and 150 μm in B, up, right.

Image analysis of Ki67 immunolabeling showed small areas enriched with Ki67+ newly-proliferated cells in at least two types of proliferative spatial arrangement (Figure 2A). The first type consisted of up to two KI67+ newly-proliferated cells named single, and a second type with multiple cells (more than three cells) called cluster. We used connected components analysis as a method to identify and quantify connections between Ki67+ newly-proliferated cells as well as to classify their arrangements. Assessing cell arrangement allows the identification of proliferative areas. Quantitative analysis in the ipsilateral region showed that most Ki67+ newly-proliferated cells presented a single arrangement (up to two cells). A fifth part presented a clustered arrangement (Figure 2B).

**Figure 2.**
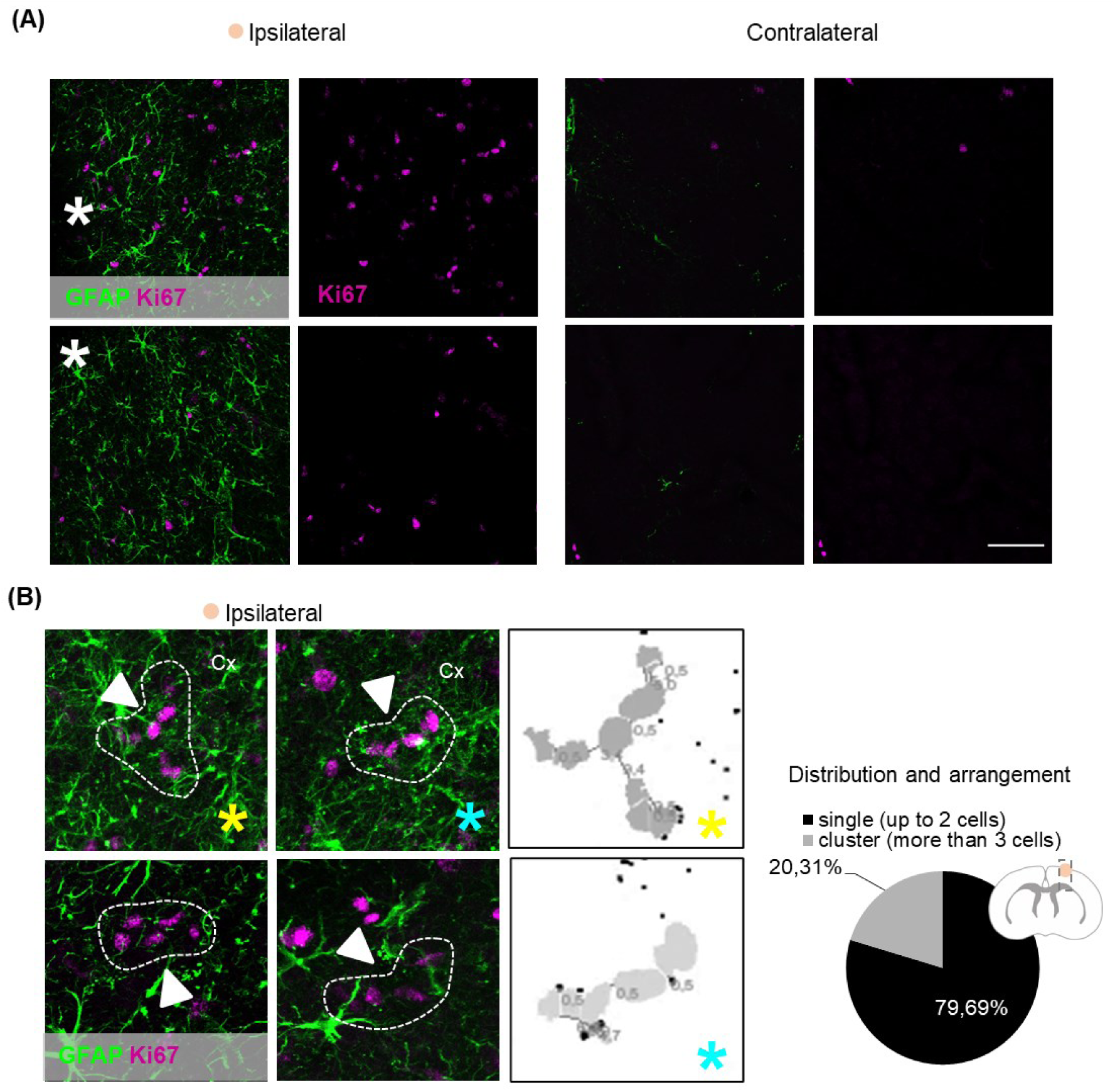
Distribution and arrangement of newly-proliferated cells. (A). Representative images of the distribution and arrangement of newly proliferated cells in the ipsilateral and contralateral cortex. We noted the appearance of many Ki67 newly-proliferated cells in cluster arrangements that resemble the configuration found in the neurogenic niches. (B). Connected component analysis of the Ki67+ cells in the ipsilateral cortex showed that at least a fifth of the cells had cluster configuration. *White, injury site; *yellow and cyan, connected component reconstruction, cells joining a cluster had a distance less than 10µm between each other. Values with p < 0.05 were considered statistically significant. Scale bar 50 μm in B, up and 20 μm in B, down.

We next analyzed the activation of Notch, Shh and Wnt signaling pathways. Neurogenic signaling markers used were neurogenic locus Notch homolog protein 1, also known as Notch1 intracellular domain (NICD, the active domain of Notch receptor), active β-catenin (β-catenin) to identify Wnt signaling activation, and Glioma-associated oncogene or Zinc finger protein (Gli1) for Shh signaling activation. As mentioned, Notch signaling activation causes the release of the cytoplasmic portion of Notch receptor (NICD). NICD binds the transcription factors RBPJk/MAML to activate Notch target genes. In parallel, Wnt signaling activation causes the translocation of cytoplasmic β-catenin into the nucleus to bind the transcription factor complex TCF/LEF and activate Wnt target genes. Shh signaling activation causes Gli1 translocation to the nucleus, acting as a transcription factor in Shh target genes (Urbán et Guillemont, 2014; Choe et al., 2015, Figure 3A). We used an immunolabeling visualization strategy based on pseudocolor image Look-Up table (LUT) combined with radial grid patterns to differentiate high intensity-labeling areas and determine the presence and distribution (distance) of the markers of neurogenic signaling activation. As mentioned, the LUT transformation creates a colorized image where gray values intensities match red, green, and blue (RGB) values. The radial grid pattern started in the lesiońs borders, and each line of the grid was 100µm apart (Figure 3B). Quantitative analysis surprisingly showed distinct signaling activation-intensities. Notch and Wnt signaling proteins NICD and (active) β-catenin showed large areas of high-intensity immunolabeling, reaching equal distances in the ipsilateral cortex (NICD, 866.6 +/− 229.7 μm; β-catenin, 785.7 +/− 299.01 μm). Nevertheless, Shh signaling protein Gli1 showed poor immunolabeling and distribution. Intensity values of each neurogenic pathway let us to establish two regions of signaling activation. A high intensity-labeling signaling region in red, named core, and a low intensity-labeling signaling region in blue, named periphery. Of note, Notch and Wnt signaling proteins had a pronounced, high intensity-labeling core (NICD, 511.100 +/− 117.1 μm; β-catenin, 400+/−230.9 μm), and a shorter low intensity-labeling periphery (NICD 355.6 +/− 113 μm; β-catenin, 385.7 +/− 69.01 μm) (Figure 3C).

**Figure 3.**
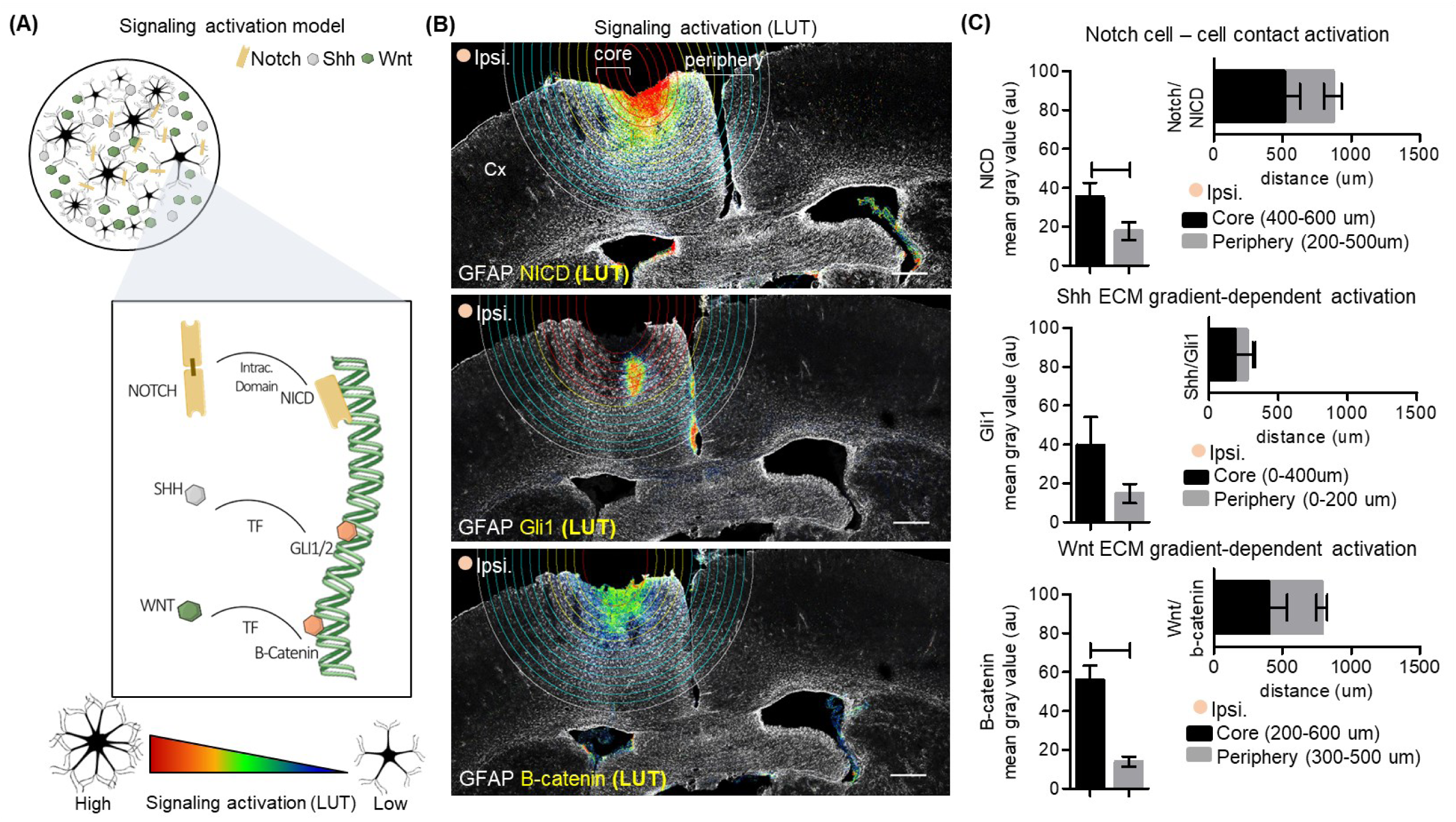
Neurogenic signaling activation. (A) Neurogenic signaling activation model. (B) Representative images of GFAP (8 bits) and NICD (Notch/LUT), Gli1 (Shh/LUT), B-catenin (Wnt/LUT) immunolabeling, and radial grid patterns. (C). Quantitative analysis of the intensity of NICD (Notch), Gli1 (Shh), and B-catenin (Wnt) immunolabeling and the distribution of signaling regions (core and periphery). The radial grid pattern started in the lesiońs borders, and each line of the grid was 100µm apart. Notch and Wnt signaling proteins, NICD and (active) β-catenin showed large areas of high-intensity immunolabeling, reaching equal distances in the ipsilateral cortex. Nevertheless, Shh signaling protein Gli1 showed poor immunolabeling and distribution. Values with p < 0.05 were considered statistically significant. Scale bar 200 μm.

### Reactive astrocytes had an aggregation response pattern in the core of Notch signaling

Given these results, we questioned whether neurogenic signaling distribution might support complex interactions that affect the network’s response to brain injury. Specifically, we asked whether GFAP reactive astrocytes present different response patterns according to their position either in the core or the periphery of Notch signaling. We analyzed GFAP protein immunolabeling and nearest neighbor distance ratio (NND ratio, Figure 4A). Reactive astrocytes in the core and the periphery of Notch signaling showed a high variability (standard deviation) in GFAP immunolabeling (mean gray value). The NND ratio of GFAP reactive astrocytes in the core (1.12 +/− 0.07) was lower compared to the periphery (1.85+/− 0.02). As mentioned, NND is a descriptor of the pattern of distribution of a population, GFAP reactive astrocytes with lower NND ratio indicates an aggregation distribution. Additional measures such as density and expected distance did not show any difference (Figure 4B).

**Figure 4.**
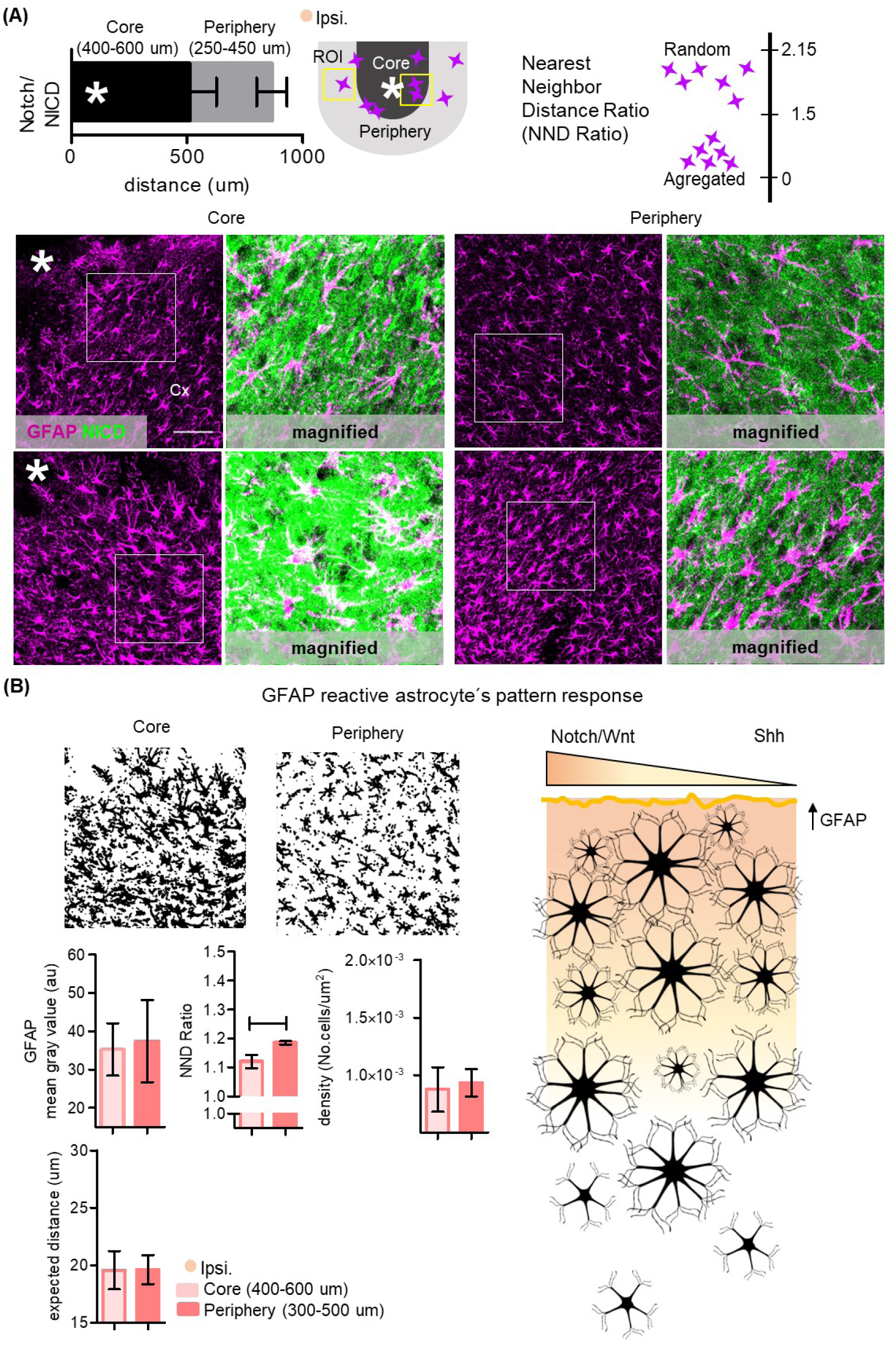
Reactive astrocytés distribution in Notch signaling regions. (A). We selected regions of interest (ROIs) in the core and the periphery of Notch (NICD) signaling. We used the Nearest Neighbor Distance Ratio (NND Ratio) to determine the reactive astrocytés response pattern (distribution). Lower ratios describe an aggregatiońs response. In contrast, higher ratios correspond with a random distribution. Representative images of GFAP/NICD immunolabeling at the core and the periphery. *Site of the lesion. (B). Quantitative analysis of GFAP, NND Ratio, density, and distance between cells. Right. Schematic representation of Notch, Wnt, and Shh neurogenic signaling and reactive astrocytes response. We found that reactive astrocytes had an aggregated response pattern in the core of Notch signaling. Values with p <0.05 were considered statistically significant. Scale bar 50 μm.

### In vitro regulation of Notch neurogenic signaling in reactive astrocytes activate neural precursors mechanisms

Findings led us to investigate the effect of Notch signaling inhibition in the transcriptional program of reactive astrocytes. We used an *in vitro* model of wound healing and reactivation by scratch. Control groups consisted of naïve astrocytes. First, we verified astrocyte response to our model of wound healing. Three days post-assay (3dpa) we performed immunocytochemical analysis of astrocyte reactivation response. Reactive astrocytes underwent cytoskeletal reorganization showing enlarged cell bodies, and hypertrophic primary and secondary branches that extended toward the injury site. Quantitative analysis showed increased GFAP immunolabeling (mean gray value). There were no differences in the number of GFAP cells, however there was a tendency to a proliferative response (%Ki67+ cells, when compared with the control group, naïve astrocytes, Figure 5A). Gene expression analysis (fold change, FC) of genes of interest, GOIs, showed the differences between control and reactive astrocytes. Some GOIs showed large inter-individual variability. For instance, in the control group GFAP showed a coefficient of variation (CV) of 48.01%, while NESTIN had a CV of 65.54%. Similar results were obtained in the scratch group, where GFAP showed 32.86%, while NESTIN showed 49.23%.

**Figure 5.**
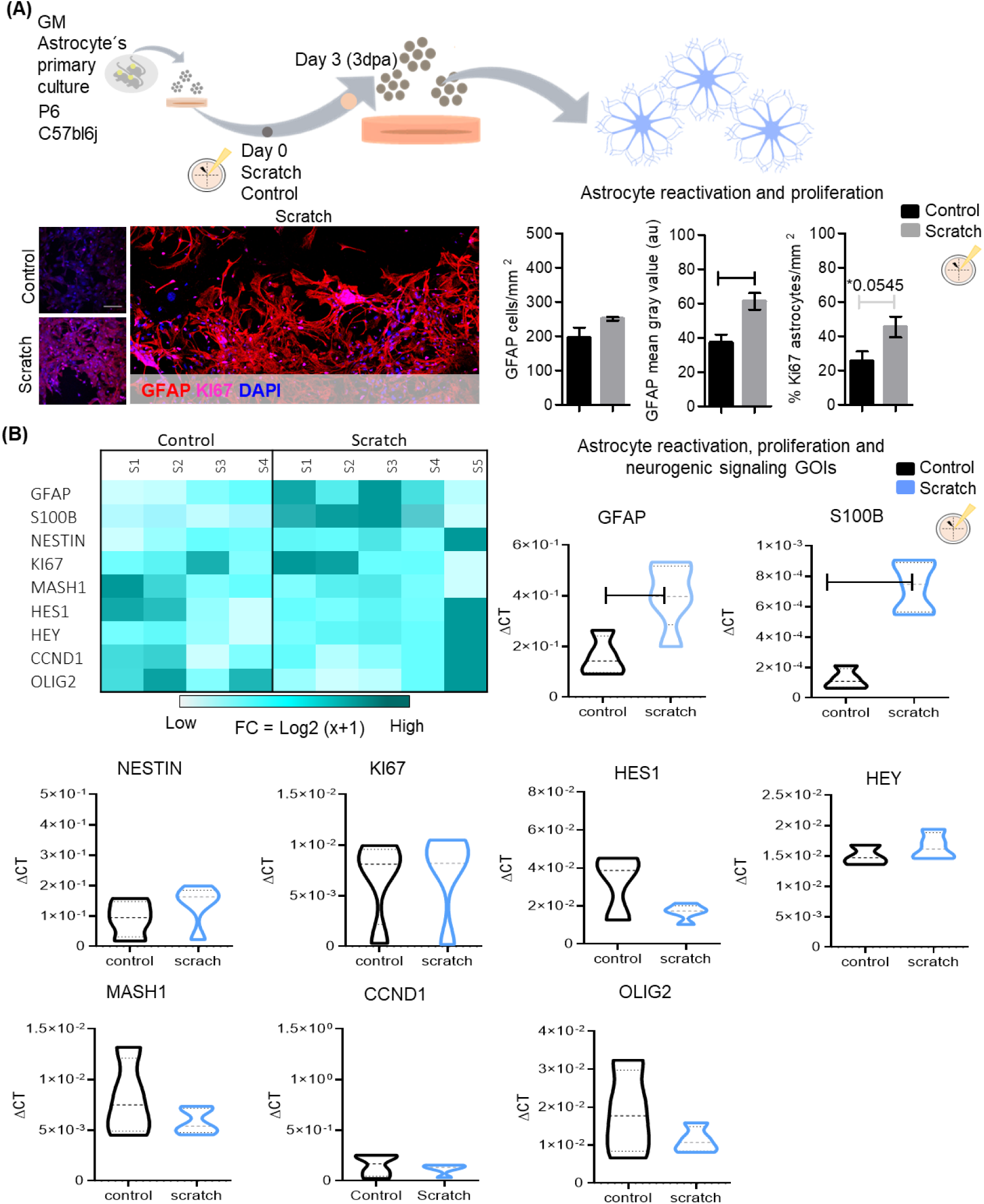
*In vitro* model of wound healing and astrocyte reactivation (A). Methodological design. Primary cultures of astrocytes at the second passage were submitted to scratch assay. Three days post-assay (3dpa) were analyzed astrocyte reactivation, proliferative response, and neurogenic signaling activation. Representative images of GFAP/Ki67+ proliferative reactive astrocytes. Branches of reactive astrocytes frequently overlap in a projection to the lesion core. Note the presence of GFAP/Ki67+ proliferative cells on the border of the lesion. Quantitative analysis of astrocyte reactivation. Number of GFAP cells, GFAP immunolabeling and proliferative response, %KI67 astrocytes. (B). Fold change-heatmap of genes of interest, GOIs. We found a higher variability between the samples. Accordingly, with the immunocytochemical analysis, reactive astrocytes showed increased expression of the genes of reactivation response GFAP and S100β. There were no changes in proliferative and reactive response genes, KI67 and NESTIN, Notch signaling genes, HES1 and HEY, Wnt-Shh signaling gene, CCND1, and neural differentiation Basic-Helix-Loop-Helix, BHLH gene, MASH1. There was an expected slight increase in HEY expression. Fold change (FC) corresponded to *Log2* (*x* + 1). N= 4 biological replicates for control group and 5 biological replicates for scratch group. We considered three technical replicates for each GOI. Values with p < 0.05 were considered statistically significant. Scale bar 50 μm.

In accordance with the immunocytochemical analysis, reactive astrocytes showed increased expression of the genes of reactivation response GFAP and S100β compared with control (naïve astrocytes). There were not changes in proliferative and reactive response genes, KI67 and NESTIN, Notch signaling genes, HES1 and HEY, Wnt-Shh signaling gene, CCND1, and neural differentiation Basic-Helix-Loop-Helix, BHLH gene, MASH1. Although there was an expected slight increase in HEY expression (Figure 5B).

In the sequence, we assessed the effect of Notch signaling inhibition in reactive astrocyte response. We considered the following groups: control reactive astrocytes, and reactive astrocytes treated with LY450139 Semagacestat, Notch signaling inhibitor. We also included an additional control group for LY450139 Semagacestat’s vehicle (0.001% DMSO). LY450139 Semagacestat is a chemical inhibitor of the γ-Secretase enzyme. The γ-Secretase enzyme participates in wide variety of biological functions, such as the formation of amyloid-β peptides and Notch signaling activation (Struhl et Greenwald, 1999, Kopan et Ilagan., 2004, Choe et al., 2015). In the case of Notch signaling, after ligand binding, the γ-Secretase enzyme cleaves the cytoplasmic portion of the Notch1 receptor releasing the Notch active domain (NICD), which translocates into the nucleus (Choe et al., 2015). A previous work demonstrated that LY450139 Semagacestat inhibited the active form of Notch in H4 human glioma in a dose dependent-manner (Mitani et al., 2012). The study showed that after 10nM dose, the activity of the active Notch (NICD) reporter dramatically decreased and showed a total inhibition at 1µM and 10µM dose (IC50=14.1 nM). Of note, the treatment would only be effective on reactive astrocytes actively signaling through Notch pathway. Notch inhibition (1μM LY450139 Semagacestat) markedly reduced reactive astrocyte genes, GFAP and S100β, but increased NESTIN. Notch signaling target genes showed a paradox response. HEY gene expression was abolished while we found an increased expression of HES1 gene and decreased MASH1 gene. Notch inhibition also increased the expression of the genes of proliferative response and Wnt-Shh signaling effectors, CCND1 and OLIG2 (Figure 6A). LY450139 Semagacestat’s vehicle (0.001% DMSO) did not show any effect in reactive response genes (Figure 6B).

**Figure 6.**
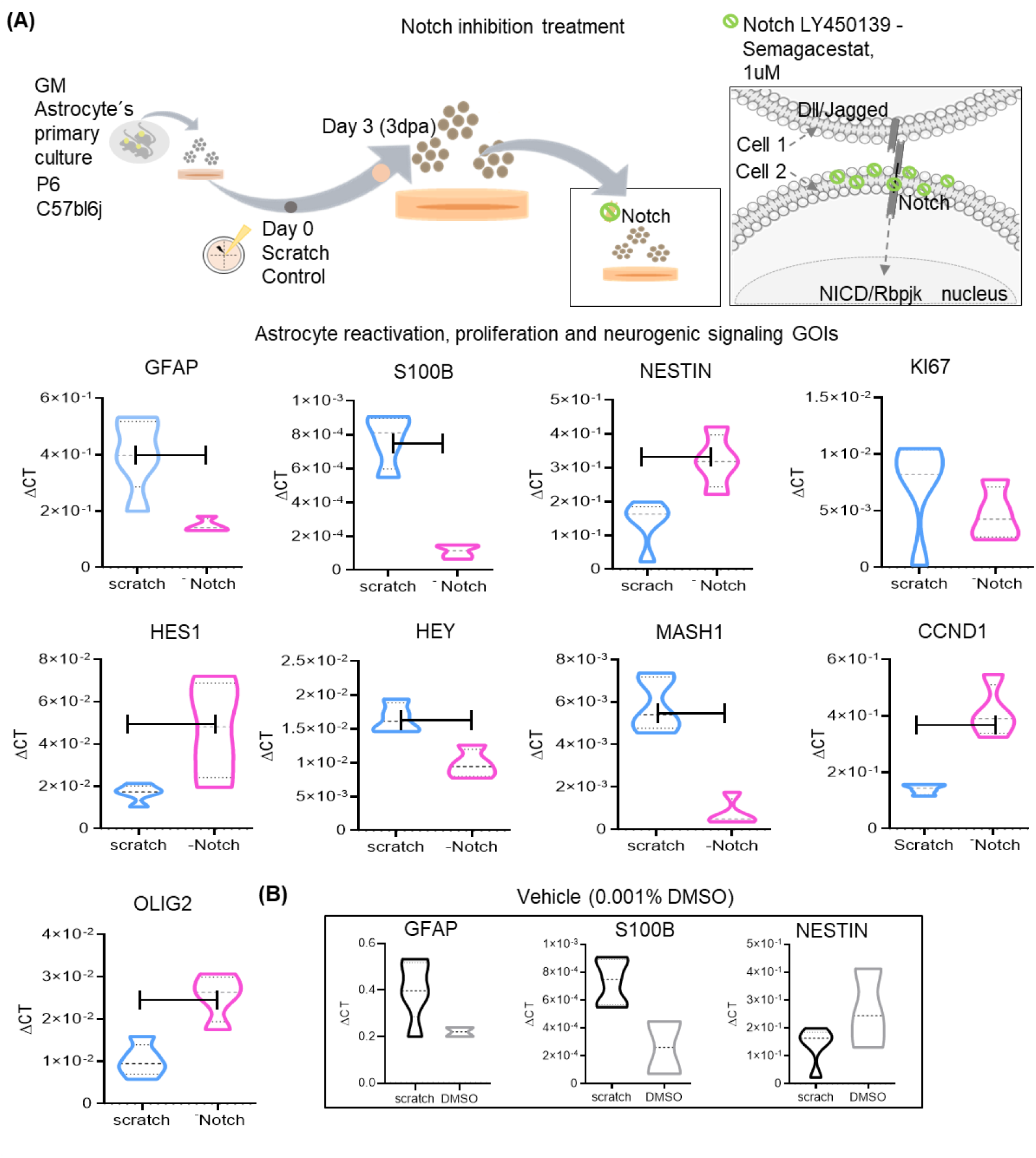
Notch inhibitory treatment in reactive astrocyte response. (A). Experimental design, schematic representation of Notch signaling inhibition and GOIs expression. Three days post-assay (3dpa) reactive astrocytes were treated with LY450139 Semagacestat, the Notch signaling inhibitor (-Notch). Reactive astrocytes genes, GFAP and S100B, Notch signaling effectors HEY, and MASH1 genes expression were abolished while we found an increased expression of NESTIN and HES1 gene. Notch inhibition also increased the expression of the genes of proliferative response and Wnt-Shh signaling effectors, CCND1 and OLIG2. (B). Semagacestat’s vehicle (0.001% DMSO) did not show any effect in reactive response genes. Values with p < 0.05 were considered statistically significant. N= 5 biological replicates for scratch group and 4 biological replicates for Notch inhibition group (LY450139 Semagacestat). We considered three technical replicates for each GOI.

### Notch signaling inhibition promoted proliferation of reactive astrocyte-derived neurospheres

As a second approach to investigate the role of Notch inhibition in reactive astrocytés proliferative response, scratched-reactivated astrocytes were seeded in low attachment surface with conditioned medium enriched with growth factors to generate self-renewing and multipotent neurospheres. Neurospheres were treated daily with LY450139 Semagacestat, Notch signaling inhibitor. We tested five working concentrations: 10 nM, 30 nM, 100 nM, 300 nM and 1 µM. We also included an additional control group for LY450139 Semagacestat’s vehicle (0.001% DMSO). After one week-treatment, neurospheres were imaged for area size calculation (Figure 7A). We found a dose-dependent effect on Notch inhibition by LY450139 Semagacestat over neurosphere size. Pairwise comparison revealed that neurosphere area of 10nM group was significantly larger (Figure 7B). Considering the large variability of the sample, we next classified the neurospheres according with the size and analyzed their frequency on each group. We identified three neurospherés sizes: small (62.79µm^2^ to 169.45 µm^2^), medium (169.46 µm^2^ to 276.10 µm^2^) and large (276.11 µm^2^ to 382.75 µm^2^). The analysis showed vast majority of neurospheres in all groups were classified as small and medium. In the control group (without LY450139 Semagacestat) 72.09% neurospheres were classified as small, 25.58% medium and 2.32% large size. In contrast, in the 10nM group, 48.65% neurospheres had small size, 45.95% were medium and 5.4% were large. Proportion of medium sized neurospheres was higher in the 10nM group. Interestingly, we found increased number of large sizes neurospheres in the groups 10nM (5,4%) and 1nM (5%). Even though the difference in neurosphere area was modest, our results suggest that Notch inhibition promoted self-renewal and proliferation of reactive astrocytes-derived neurospheres (Figure 7C).

**Figure 7.**
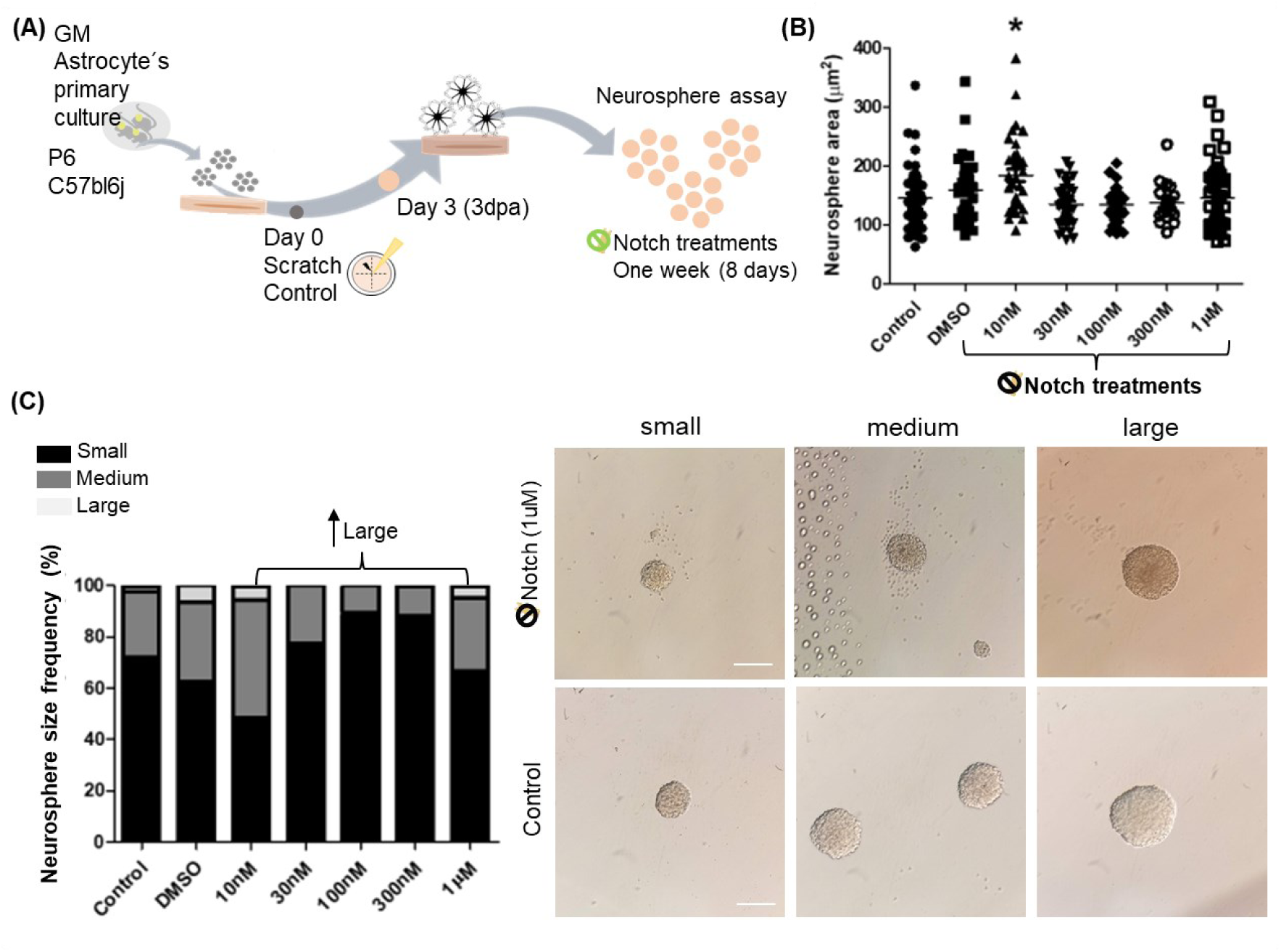
Notch signaling inhibition in reactive astrocyte-derived neurospheres. (A) Methodological design. Primary culture of astrocytes at second passage were submitted to scratch assay. Three days post-assay (3dpa) reactive astrocytes were seeded at a cell density of 8×10^3^ cells/well in a 24-well plate with low attachment surface (poly-2-hydroxyethyl methacrylate). Cells were fed with conditioned medium enriched with growth factors (20 ng/ml bFGF, and 10ng/ml EGF). Neurospheres were treated daily, for one week (8 days), with LY450139 Semagacestat. (B). We tested five working concentrations (groups) (10 nM, 30 nM, 100 nM, 300 nM and 1 µM). We also included an additional control group for LY450139 Semagacestat’s vehicle (0.001% DMSO). Neurosphere area size quantification analysis. (C). Neurosphere size frequency. The relative number of neurospheres in each group is expressed as a percentage of the total number of neurospheres. Representative images of small, medium, and large size neurospheres. We found a dose-dependent effect on Notch inhibition by LY450139 Semagacestat over neurosphere size. The analysis showed vast majority of neurospheres in all groups were classified as small and medium. We found increased number of large sizes neurospheres in the groups 10nM (5,4%) and 1nM (5%). Even though the difference in neurosphere area was modest, our results suggest that Notch inhibition promoted self-renewal and proliferation of reactive astrocytes neurospheres. Values with p < 0.05 were considered statistically significant. Scale bar 20 μm.

### Crosstalk between Notch inhibition and Wnt-Shh signaling in reactive astrocytes profile

Since the activation of specific signaling pathway is essential for reactive astrocyte neuroprotective response to brain injury, we assessed the effect of Wnt-Shh signaling (0,25 ug/mL rmShh, and 4ug/mL rmWnt-3a) in reactive astrocytes treated with Notch signaling inhibitor (1µM, γ-Secretase inhibitor LY450139 Semagacestat). We considered the following groups: reactive astrocytes; reactive astrocytes treated with LY450139 Semagacestat, Notch signaling inhibitor; and reactive astrocytes treated with LY450139 Semagacestat, rmShh, and rmWnt-3a. (Figure 8A). Gene expression analysis showed downregulation of reactive astrocytes genes GFAP and S100β, and increased expression of the genes of proliferation and Wnt-Shh signaling effector, CCND1, and neural differentiation, OLIG2, when compared with reactive astrocytes. HES1, Notch signaling effector, was markedly downregulated. HEY expression persisted downregulated. MASH1 gene, did not show differential expression. Results suggest effective Notch signaling inhibition (HES1 and HEY, Figure 8B).

**Figure 8.**
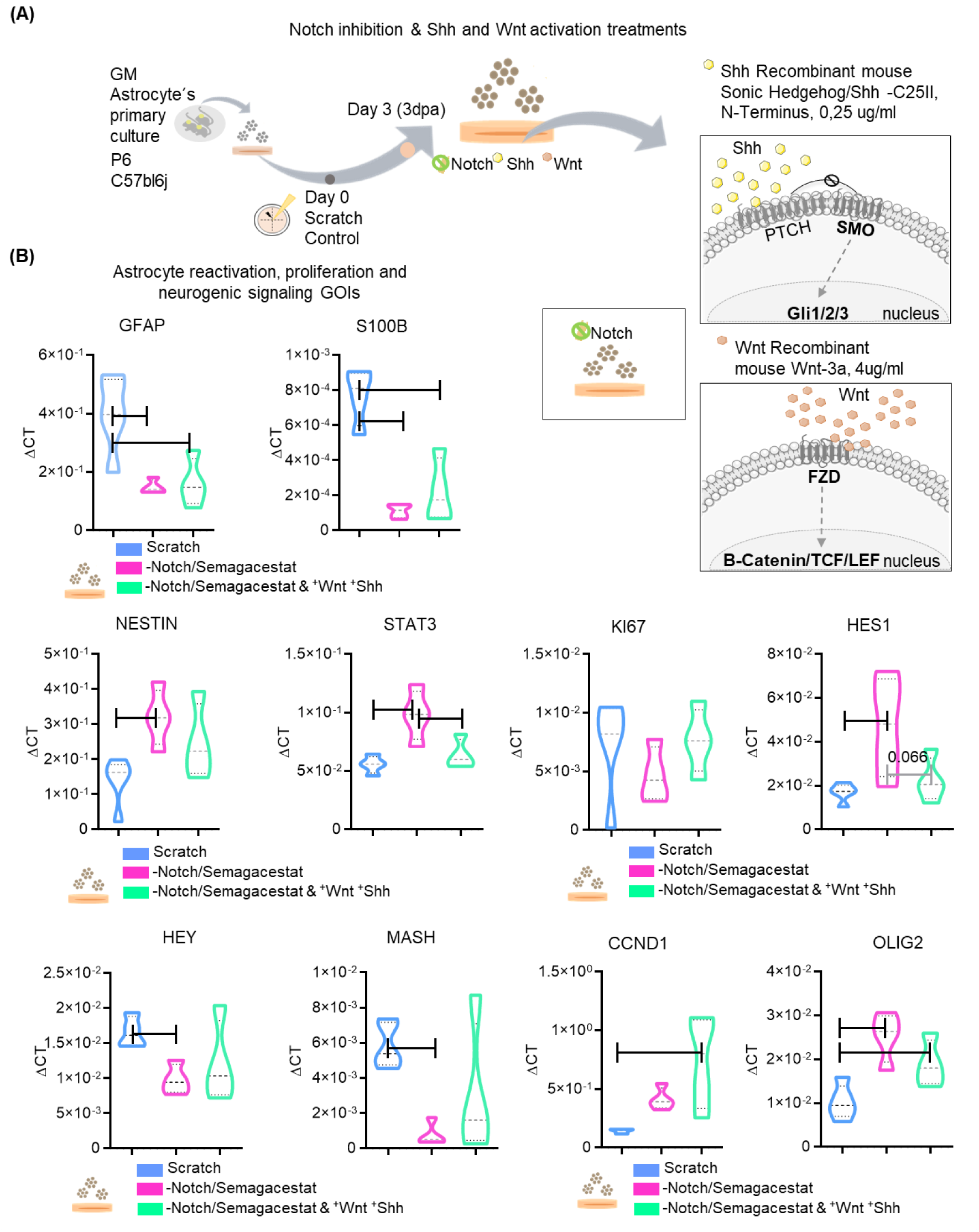
Notch signaling inhibition and Wnt-Shh activation treatment. (A). Experimental design and schematic representation of Wnt/Shh (rmShh/rmWnt-3a) signaling. Three days post-assay (3dpa) reactive astrocytes were treated with LY450139 Semagacestat, the Notch signaling inhibitor (-Notch/Semagacestat), and rmShh/rmWnt-3a (-Notch/Semagacestat & +Wnt+Shh). We analyzed astrocyte reactivation, proliferative response and neurogenic signaling. (B). GOIs expression. Analysis showed downregulation of reactive astrocytes genes, GFAP and S100β, and increased expression of CCND1 and OLIG2, Wnt-Shh signaling effectors. HES1 gene was markedly downregulated. Results suggest effective Notch signaling inhibition. N= 5 biological replicates for scratch group, 4 biological replicates for Notch inhibition group (LY450139 Semagacestat) and 4 biological replicates for Notch inhibition & +Shh+Wnt group. Values with p < 0.05 were considered statistically significant. We considered three technical replicates for each GOI.

## Discussion

A signal produced by a defined source forms a concentration gradient while it spreads through the tissue, allowing intracellular biochemical cascades for gene expression. Traumatic brain injury (TBI) creates a temporal microenvironment activating specialized cellular programs for repair and regeneration. Neurogenic signaling pathways modulate neural progenitoŕs proliferation, migration, and cell fate decisions. Likewise, reactive astrocytes trigger a transcriptional-proliferative program where neurogenic signaling molecules play crucial roles. Precise molecular mechanisms are context-specific and are not fully understood. We studied cellular and molecular aspects of TBI-reactive astrocyte response after Notch-Wnt neurogenic signaling modulation. The model of brain damage by TBI creates an extensive area of tissue loss in the somatosensorial cortex that resembles severe conditions such as penetrating injuries. Astrocytes will acquire a reactive-proliferative program that creates a non-reversible glial scar. Accordingly, we found strong reactive astrocyte response in the ipsilateral region. Reactive astrocytes showed enlarged cell bodies and upregulation of GFAP and S100B proteins. There was a progressive increase in Ki67+ newly-proliferative cells, distributed almost exclusively near the injury site. Half of newly-proliferative population were GFAP reactive astrocytes (%Ki67/GFAP). Most newly proliferated cells presented a single arrangement (up to two cells), and just a fifth part exhibited a clustered arrangement. Neurogenic signaling analysis showed Notch (NICD) and Wnt (active β-catenin) regional upregulation and signaling concentration gradient in the ipsilateral cortex.

Models of brain damage include a wide variety of stimuli. For instance, brain damage and neuroinflammation models used for assessing neurogenic signaling response have included cryo-injury (Tatsumi et al., 2010), stroke/ischemia/MCAO (Shimada et al., 2011; Mastroiacovo et al., 2009; Magnusson, 2014; LeComte et al., 2015; Zhong et al., 2018), LPS (Acaz-Fonseca et al., 2019) and controlled cortical impact (Salehi et al., 2018). Given this variety of stimuli, it is not surprising the appearance of conflicting findings. The type of stimulus, severity, and anatomical localization/astrocyte subpopulation might produce a differential reactivation response (Burda & Sofroniew, 2014). Likewise, there is no consensus on the role of Notch, Wnt, and Shh signaling in reactive astrocyte function. Some studies suggest that reduced Notch signaling is necessary for reactive astrocyte neuroprotective response (Magnusson, 2014; Acaz-Fonseca et al., 2019), while others have found that Notch signaling activation is essential for astrocytés proliferative reactive response (Shimada et al., 2011; LeComte et al., 2015; Zhang et al., 2015; Zhong et al., 2018). Accordingly with our findings, a previous study showed that scratched-reactive astrocytes close to the lesion core (*in vitro*) activated the Notch1/Jagged1 signaling pathway and upregulated Hes5 expression, a Notch target gene (Ribeiro et al., 2021). Moreover, Wnt signaling inhibition has been reported after ischemia/MCAO (Mastroiacovo et al., 2009) and upregulated in controlled cortical impact (Salehi et al., 2018) models. This last work showed that β-catenin upregulation in the vessels and the perilesional area at days one and seven post-injury (1-7dpi) promoted angiogenesis and vascular repair (Salehi et al., 2018).

In this context, comparison between brain damage models became challenging. Moreover, we only found one study on Notch-Wnt signaling in brain injury. The study showed that Wnt3a and β-catenin expression increased significantly seven days post-injury (7dpi) in a model of brain injury in rats, while Notch1 and Hes1 expression increased gradually over time (Wei et al., 2020). The authors suggested a synergistic effect of both pathways on repairing mechanisms in brain injury. On the other hand, our analyses also showed that Shh (Gli1) signaling was poorly detected. Gli1 is one of the three main protein effectors (Gli1, Gli2, and Gli3) in Shh signaling. We cannot reject the possibility of Shh signaling activation, since we just analyzed Gli1. Shh upregulation has been found in models of cryo-injury (Amankulor et al., 2009), stab wound injury, and ischemia (Sirko et al., 2013). In all of them, where was sufficient to activate proliferation and neuroprotective response. In opposition, other studies showed that models of brain damage by cortical impact and stab wound injury exhibited a pronounced reduction in Shh signaling (Allahyari et al., 2019; Wu et al., 2020). Altogether, our observations suggest the activation of mechanisms for the repair and renewal of the neural populations following injury.

We also described a heterogenous population of GFAP reactive astrocytes in a preferential aggregated distribution in the core of Notch signaling. Studies have shown that multiple glial cells proliferate around the margins of lost neural tissue and interact to form an astrocyte limitans border that delineates viable neural tissue from stromal cell scar tissue (Sofroniew, 2015; Wahane et Sofroniew, 2021). Moreover, organism models of stroke and intracerebral hemorrhage concluded that Notch signaling is involved in reactive astrocyte proliferation in the lesion core (or peri-infarct area, Shimada et al., 2011; LeComte et al., 2015; Zhong et al., 2018). Similarly, other study reported that the Notch1-STAT3-ETBR axis is responsible for reactive astrocyte proliferation after ischemic stroke. The authors showed that Jagged1 receptor was upregulated on the ischemic-ipsilateral region, where Notch1/Jagged1 signaling promoted reactive astrocyte proliferation (LeComte et al., 2015). Our assumption, for our brain injury model, is that some of these cells at the core of Notch signaling, in close contact with the injured region, might be *astrocyte-limitans-borders*. Indeed, we identified a discrete subpopulation of clustered, newly-proliferative cells in the ipsilateral cortex. Notch signaling might provide neurogenic support to these reactive astrocytes, while cell-cell interactions reinforce the signal.

We next assessed whether the modulation of neurogenic signaling pathways might change the gene expression profile of reactive astrocytes *in vitro*. We found that the treatment with LY450139 Semagacestat effectively inhibited Notch-HEY but not Notch-HES1 signaling. In fact, HES1 expression did not varied upon control and reactive astrocytes. Acknowledging that interactions between transcription factors are countless, complex to consider through solely gene expression analysis, and context-dependent, we propose the occurrence of at least two mechanisms by which Notch inhibition with LY450139 Semagacestat attenuated reactive astrocytes response (GFAP and S100β) and activated parallel neural precursor mechanisms (proliferation and self-renewal). The first mechanism is directly related to Notch-HEY inhibition. The second mechanism involves Notch-independent HES1 upregulation and repression of neural differentiation gene MASH1.

Thus, as a second approach, to assess whether Notch inhibition might increase the number of committed-proliferative astrocytes and activate neural precursor mechanisms, we analyzed reactive astrocytes-derived neurospheres treated LY450139 Semagacestat. Specifically, we assessed the effect of Notch inhibition in reactive astrocyte’s ability to form neurospheres. Reactive astrocytes were seeded at clonal density in low attachment surface with conditioned medium enriched with growth factors and treated daily with LY450139 Semagacestat at different working concentrations. Results showed a slight increase in neurosphere area size, mainly in the 10 nM group. Higher inhibitory treatments sustained medium and large size neurosphere growth. Altogether, our observations suggest that Notch inhibition promoted proliferation and self-renewal mechanisms.

Notch-HEY signaling in astrocyte reactivation and brain injury has not been reported. However, in the developing mouse lateral ganglionic eminence, it was demonstrated that Notch1 and HEY1 are necessary to maintain slowly dividing neural progenitor cells. Interestingly, HEY1 shRNA-knockdown mice (E14.5-E16.5) showed a reduction in the fraction of SOX2 progenitor cells and increased ASCL1 (MASH1) differentiated cells. The study suggested that persistent and high levels of Hey1 expression in neural progenitor cells are responsible for undifferentiated state maintenance in the adult brain and might distinguish slow-from rapid-cycling neural progenitor cells (Harada et al., 2021). Moreover, previous studies showed that Notch signaling inhibition is required for Müller glia progenitors (MGs) response to retinal damage (Conner et al., 2014; Wan et Goldman, 2017; Sahu et al., 2021). MGs are the main glial cell of the retina and reside in intimate contact with neurons and blood vessels. Specifically, one study revealed that Notch signaling suppression generated new progenitor cells that migrate to the injury (Sahu et al., 2021). RNAseq and ATAQseq analysis of transgenic zebrafish MGs models (tuba1a GFP fish) submitted to retinal injury by needle poke showed repression of HEY and ID2B genes, and 60% of the response-genes presented changes in chromatin accessibility. In addition, GFAP/GFP MGs cells of control (uninjured) fish treated with the γ-Secretase inhibitor DAPT (Notch signaling inhibitor) showed similar changes in HEY1 and ID2B gene expression. HEY1 knockdown (HEY1-morpholino) fish model showed that MGs created an expanded response zone after the needle poke injury model (Sahu et al., 2021). Altogether, the data suggested that Notch inhibition contributed to the MG response profile. Editing the HEY1 gene resulted in an expanded zone of injury response, upregulating the number of MG proliferative cells. Considering our results, we hypothesize whether Notch-HEY inhibitory signaling might increase or expand the number of reactive astrocytes. Altogether the data suggested that Notch inhibition contributed to the MG response profile. Editing the HEY1 gene resulted in an expanded zone of injury response, upregulating the number of MG proliferative cells. Considering our results, we hypothesize whether Notch-HEY inhibitory signaling might increase or expand the number of responsive reactive astrocytes. Notch inhibition also promoted upregulation of the reactive astrocyte cytoskeletal protein NESTIN, which is responsible for cell migration (Lepekhin et al., 2001; Wilhelmsson et al., 2019).

The second mechanism involves Notch-independent HES1 upregulation and repression of neural differentiation genes such as MASH1. As mentioned, Notch signaling represses MASH1 by activating BHLH target genes such as HES1, HES5, and HEY. Previous works had shown Notch-independent HES1 regulation. Hes1 protein could bind their promoter and regulate HES1 expression in a negative feedback loop, displaying an oscillatory effect in fibroblasts (Hirata et al., 2002) and neural progenitors (Baek et al., 2006; Shimojo et al., 2008). HES1 expression could also be regulated by diverse signaling pathways such as Shh in progenitor cells of the retina (Wall et al., 2009), and kinases in endothelial cells (Curry et al., 2006). Specifically, one study showed that hypoxic conditions induced HES1 expression through Notch-dependent and -independent mechanisms and could be cell type-specific. P19 mouse embryonic carcinoma cell line submitted to hypoxia and treated with the γ-Secretase inhibitor DAPT did not show changes in the expression of Notch target genes HES1, HEY1, and HEY2. In contrast, DAPT treatment in primary mouse brain endothelial cells submitted to hypoxia downregulated HES1 expression (Zheng et al., 2017). The study also showed that in the P19 cell line, siRNAs targeting HIF-1α or ARNT significantly reduced hypoxia-dependent induction of HES1, HEY1, and HEY2 gene expression. Interestingly, another study showed that persistent and high levels of HES1 expression are required for the formation of organizing centers that regulate neuronal differentiation. Neural progenitor primary cultures derived from E11.5 mouse embryos infected with CLIG-HES1 to promote persisting HES1 expression showed less proliferation efficiency, growth rate, and increased Cyclin D1 (CCND1 gene) immunolabeling. The study suggested that persistent HES1 expression promoted a radial glial profile in the cell culture, with reduced cell proliferation and inhibition of neurogenesis (Baek et al., 2006). Altogether, these studies indicate that multiple signals and pathways can converge to regulate HES1 expression. Our results are consistent with this interpretation, showing that reactive astrocytes treated with the inhibitor of the γ-Secretase enzyme LY450139 Semagacestat, upregulated HES1 expression in a Notch-independent manner, that might promote a precursor profile, maintaining the repression of neural differentiation genes (MASH1). Hes1 might not be the signaling effector of Notch in reactive astrocytes *in vitro*, however, we could have missed important gene expression regulations due to its oscillatory pattern.

In system biology, crosstalk between signaling pathways describes the integration of signaling from multiple pathways within a network and their biological response. To be considered as a crosstalk must be filled the following criteria, a). The signal of the network should produce a response different from the one expected by the pathways, and b) the signaling pathways should be connected directly or indirectly. Direct crosstalk is established when existing specific shared components, or when a component of one pathway is modified by enzymes that are proper of another pathway. Indirect crosstalk is established by the sequential action of different pathways, especially, when the action of a pathway is necessary for the activation of a second pathway (Vert et Chory., 2011).

Here we found a positive interaction of Wnt-Shh signaling in the Notch inhibitory profile. One group comprehensively studied the interaction between Notch-inhibition and Wnt-signaling activation in the regenerative response of mice hair cell progenitors of the inner ear (Li et al., 2015; Wu et al., 2016; Ni et al., 2016; Wu et al., 2021). Moreover, their most recent publication demonstrated the crosstalk between Notch, Wnt, and Shh signaling pathways in the reparative response of the cochlea after cytotoxic neomycin treatment. The study showed that Notch inhibition and SHH-Wnt treatment promoted progenitor proliferation and activation of a regenerative response (Wu et al., 2021). The study evaluated the combinatorial effects of the inhibitor DAPT, the Wnt agonist QS11, and recombinant Sonic Hedgehog (rmShh) in mice cochlea and their potential regenerative response after neomycin treatment. The study found that the simultaneous treatment with the inhibitor of the y-secretase enzyme DAPT and Wnt agonist QS11 produced more Edu+ proliferative cells than the DAPT-only group. After cytotoxic treatment, the number of EdU + cells in the DAPT-QS11 group were greater than in the DAPT-only group. Even more, the DAPT-QS11-rmShh group generated many more EdU+cells in both the intact and damaged cochlea. A later RNAseq analysis showed that the proliferative response in the DAPT-QS11-rmShh group was mainly induced via SHH and Wnt signaling activation. Our results indicated an indirect crosstalk of Wnt-Shh signaling pathways in the regulation of BHLH and Notch target genes HES1 and HEY after Notch inhibition. In addition, Notch inhibition might also have a supportive effect on the Wnt-Shh target gene CCND1 and OLIG2.

CCND1 is the gene that encodes the protein cyclin D1, a well-known G1 cell-cycle regulator that also might have a dual role in progenitor cells in a cell-cycle independent manner (Fu et al., 2004; Lukaszewicz et Anderson, 2011; Mao et al., 2019). One study showed that Cyclin D1 promoted neurogenesis in the chick spinal cord. Image analysis showed that Cyclin D1 was persistently expressed in motoneuron progenitors of the developing spinal cord during the initial phase of differentiation. Experiments using siRNA targeting Cyclin D1 and Cyclin D2 expression showed that just Cyclin D1 promoted *in vivo* neurogenesis. Moreover, the study found a possible co-interaction with HES6, a Notch signaling repressor (Lukaszewicz et Anderson, 2011). Another study also found that overexpression of CCND1 in retinal organoids promoted neurogenesis. The study showed that in progenitor cells, CCND1 was expressed in a cell-cycle-independent manner. Using an inducible system to create an ectopic overexpression they found that CCND1 increased the number of neuronal cells in retinal organoids. The results indicated that CCND1 overexpression induced more cells to leave the cell cycle and commit to neuronal fate via increased ASCL1 (also known as MASH1) expression (Mao et al., 2019). OLIG2 is a BHLH transcription factor and Shh target gene that promotes glial differentiation. OLIG2 is highly expressed during CNS development (Lu et al., 2000, Zhou et al., 2002). One study used an *in vitro* model of mice embryonic stem cells exposed to a neural progenitor induction medium and evaluated the transcriptional changes in the transition of neural progenitors to motor neuron progenitors. The study showed that treatment with Smoothened/Shh signaling agonist (SAG) resulted in the generation of neural progenitors that expressed markers of ventral spinal cord development such as OLIG2 and NKX2.2. Single-cell RNAseq analysis showed two functions of OLIG2 as developmental regulator. OLIG2 established motor neuron progenitor identity downstream of Shh signaling activation and promoted neuronal differentiation in motor neuron progenitors by suppressing the expression of Notch target genes HES1 and HES5 (Sagner et al., 2018). We cannot directly assume the role of CCND1 in our model of astrocyte reactivation. However, since we found that reactive astrocytes activated neural precursors mechanisms after Notch inhibition, we suggest that combined Wnt-Shh treatment might promote proliferation and neural differentiation via CCND1 and OLIG2 upregulation, and HES1 and HEY downregulation.

## Conclusion

Our results provide new evidence of cortical Notch-Wnt signaling activation after TBI. Reactive astrocytes in the core of Notch signaling showed a differential aggregated distribution. *In vitro*, Notch inhibition promoted a neural precursor profile and might increase the number of cells committed in a proliferative response. Finally, we found an indirect co-regulation of Wnt-Shh signaling in BHLH-Notch target genes and a Notch-supportive effect in Wnt-Shh signaling activation.

## Author Contributions

Conceptualization, LMDG and MP; methodology and analysis, LMDG, TNR, JCB, NRC; data curation, LMDG and MP; writing-original draft preparation, LMDG, TNR, JCB, NRC, MP; writing-review and editing, LMDG, JCB and MP; visualization, LMDG; supervision, MP; funding acquisition, MP. All authors have read and agreed to the published version of the manuscript.

## Institutional Review Board Statement

The animal study protocol was approved by the Ethics Committee of Universidade Federal de Sao Paulo (CEUA codes 1364051015, 2451111116, and 7740290318 and date of approval).

## Acknowledgments

We are thankful to Priscila Nicolicht, PhD, Amanda Arnaut, MSc and Agnes Sardinha Pinto, MSc for assistance in the experiments. We are also grateful for the technical support of the Confocal Microscopy Unit, INFAR/UNIFESP, and the Animal Facility CEDEME/UNIFESP.

## Materials and Methods

### Animals

Animals were handled in compliance with the current Brazilian legislation regarding the care and use of experimental animals. Procedures were approved by the correspondent Committee on Ethics in the Use of Animals of the Federal University of Sao Paulo, UNIFESP (CEUA No. 1364051015, 2451111116, and 7740290318). CEDEME/UNIFESP (Sao Paulo, Brazil) animal facility supplied isogenic C57Bl/6J mice, aged 1 and 45 days (P1 and P45). The animals were housed in standard cages, maintained under controlled light-dark cycles (12hs-12hs) with food and water ad libitum. We made all efforts to minimize suffering and the number of animals used.

### Model of brain damage by traumatic brain injury (TBI)

We used a model of brain damage by traumatic brain injury (TBI) previously described in Mundim et al. (2019). Briefly, adult (P45), male, C57BL/6j mice were deeply anesthetized with intraperitoneal injection of 0.2% acepromazine (2.5 mg/kg, Vetnil, Louveira, SP, Brazil), 2% xylazine (20 mg/kg, Syntec, SP, Brazil), 10% ketamine hydrochloride (80 mg/kg, Syntec, SP, Brazil), and 0.05% fentanyl (0.5 mg/kg, Syntec, SP, Brazil). After conscious and pain assessment, mice were placed in a stereotaxic frame, shaved on the top of the head, and submitted to a unilateral penetrating lesion performed with a 22-gauge needle (0.7 mm) in the left somatosensorial cortex with the following stereotaxic coordinates (relative to Bregma), anteroposterior, AP 0 mm, medial-lateral, ML 1 mm, dorsoventral, DV − 0.7 mm. The lesion was performed three times for 2 minutes. Postoperative care included controlled temperature at 37°C and 0.05% fentanyl (0.5 mg/kg, Syntec, SP, Brazil). Mice were left to recover on a heating pad until they were fully awake. For proliferation analyses, mice received a daily injection of BrdU (75mg/kg, Sigma Aldrich, St. Louis, USA). Three days post-lesion (3dpl), mice were deeply anesthetized with intraperitoneal injection of 2% xylazine, 10% ketamine hydrochloride, and 0.05% fentanyl and intracardially perfused with 4% paraformaldehyde (PFA). Brains were removed from the skull, fixed in 4% PFA overnight at 4°C, submersed in 30% sucrose overnight at 4°C, and frozen using dry ice. Cryostat coronal sections of 40-50µm were collected and prepared for immunohistochemistry.

Astrocyte primary culture was adapted from Yang et al. (2009). Briefly, neonatal (P1), C57BL/6j mice were deeply anesthetized (10% ketamine hydrochloride 100 mg/kg, Syntec, SP, Brazil, and 2% xylazine, 10 mg/kg, Syntec, SP, Brazil) and euthanized by decapitation. Brain cortices were carefully dissected and placed in HBSS solution (5.36mM KCl, 0.44mM KH2PO4, 4.16 mM NaHCO_3_, 136.9 mM NaCl, 0.336 mM Na_2_HPO_4_, 5.55 mM Glucose in distillated H_2_O, Ca^2+^ and Mg^2+^ free). Meninges were carefully removed, and tissue was mechanically dissociated and incubated with 0.25% trypsin (Sigma Aldrich, St. Louis, USA) in Versene solution (2.7 mM KCl, 1.8 mM KH_2_PO_4_, 137 mM NaCl, 10 mM Na_2_PO_4_, 0.68 mM EDTA in distillated H_2_O) at 37°C for 20 minutes. Sequentially, trypsin activity was blocked with the addition of fetal bovine serum (FBS) (1:2, Cultilab, Campinas, Brazil). The cells were transferred to 15ml tubes and centrifuged at 200g for 5 minutes and were resuspended in Versene solution. Cells were dissociated and homogenized using different pipette tips size and filtered through a 40 μm cell strainer. Finally, dissociated cells were resuspended in astrocyte medium DMEM/F12 containing 10% FBS, 1% L-glutamine (200mM, Sigma Aldrich, St. Louis, USA) and 1% penicillin/streptomycin (100 IU, Gibco, Grand Island, USA). Finally, the cells were plated in T25 flasks coated with poly-l-lysine (Sigma Aldrich, St. Louis, USA) and incubated at 37°C in CO_2_ incubator. Half-medium changes were performed every 2-3 days.

### Cortical astrocyte primary culture

Primary culture of astrocyte was adapted from Yang et al., (2009). Briefly, neonatal (P1), C57BL/6j mice were deeply anesthetized (10% ketamine hydrochloride 100 mg/kg, Syntec, SP, Brazil, and 2% xylazine, 10 mg/kg, Syntec, SP, Brazil) and euthanized by decapitation. Brain cortices were carefully dissected and placed in HBSS solution (5.36mM KCl, 0.44mM KH_2_PO_4_, 4.16 mM NaHCO_3_, 136.9 mM NaCl, 0.336 mM Na_2_HPO_4_, 5.55mM Glucose in distillated H_2_O, Ca^2+^ and Mg^2+^ free). Meninges were carefully removed, and tissue was mechanically dissociated and incubated with 0.25% trypsin (Sigma Aldrich, St. Louis, USA) in Versene solution (2.7 mM KCl, 1.8 mM KH_2_PO_4_, 137 mM NaCl, 10 mM Na_2_PO_4_, 0.68 mM EDTA in distillated H_2_O) at 37oC for 20 minutes. Sequentially, trypsin activity was blocked with the addition of fetal bovine serum (FBS) (1:2, Cultilab, Campinas, Brazil). The cells were transferred to 15ml tubes and centrifuged at 200g for 5 minutes and were resuspended in Versene solution. Cells were dissociated and homogenized using different pipette tips size and filtered through a 40 μm cell strainer. Finally, dissociated cells were resuspended in astrocyte medium DMEM/F12 containing 10% FBS, 1% L-glutamine (200mM, Sigma Aldrich, St. Louis, USA) and 1% penicillin/streptomycin (100 IU, Gibco, Grand Island, USA). Finally, the cells were plated in T25 flasks coated with poly-l-lysine (Sigma Aldrich, St. Louis, USA) and incubated at 37°C in 95% CO^2^ incubator. Half-medium changes were performed every 2-3 days.

### Model of wound healing and astrocyte reactivation

The model of wound healing was adapted from Yang et al., (2009). We used astrocytes cultures in second passage, reaching 80-90% of confluency. Astrocytes were seeded (6-8×10^5^ cell/area) in either 13mm glass coverslips for immunocytochemistry or 34mm plate for gene expression analysis (quantitative PCR, qPCR), coated with poly-l-lysine. Carefully a 10ul pipette tip was used to scratch the surface of the culture in a graded pattern. The scratch pattern for coverslips was a “cross” composed of one horizontal and one vertical scratch, and for 34 mm dishes, the pattern was a “grid” composed of several scratches 0.5 cm distant from each other. Scratch creates areas of cell discontinuity (loss of cell-cell contact) and detachment, resembling the borders in the model of brain injury. Cells were incubated at 37°C in a 95% CO_2_ incubator. No medium change was performed after the assay. Three days post-assay (3dpa), cultures were highly enriched with reactive astrocytes and were processed for immunocytochemistry or received neurogenic signaling treatments.

### Neurogenic signaling treatments

Three days post-assay (3dpa), reactive astrocytes in 34 mm plates (6-8×10^5^ cell/area) received neurogenic signaling treatments. We followed the manufacturers’ instructions in the preparation and storage of the solutions. LY450139 Semagacestat (Santa Cruz Biotechnology, California, USA), the Notch signaling inhibitor was diluted in DMSO (Sigma Aldrich, SP, Brazil) to a stock concentration of 25 mg/mL. The working concentration used was 1µM. Recombinant mouse Sonic Hedgehog / Shh-C25II, N-Terminus (rmShh) (464-SH-025, R&D system, CA, USA) was diluted in PBS 1x and 0,1% BSA to a stock concentration of 125 µg/mL. The working concentration used was 0,25 µg/mL. Recombinant mouse Wnt-3a (rmWnt) (1324-WN-010, R&D system, CA, USA) was diluted in PBS and 0,1% BSA to a stock concentration of 40 µg/mL. The working concentration used was 4 µg/mL. In all cases aliquots were stored at −20°C. We considered the following experimental groups: control (naïve astrocytes); reactive astrocytes; reactive astrocytes treated with LY450139 Semagacestat, Notch signaling inhibitor; reactive astrocytes treated with LY450139 Semagacestat, rmShh, and rmWnt-3a; and reactive astrocytes treated with 0.001% DMSO, LY450139 Semagacestat vehicle, to eliminate vehicle interference in the assay. Cells were incubated at 37°C in 95% CO_2_ incubator and no medium change was performed after the assay. After the treatments, the cells were processed for total RNA extraction and gene expression analysis (qPCR).

### Neurosphere Assay and Notch inhibition treatment

Three days post-assay (3dpa), reactive astrocytes were washed carefully with PBS 1x and incubated at 37°C for 5 minutes with 0.25% trypsin (Sigma Aldrich, St. Louis, USA). Trypsin activity was blocked with addition of fetal bovine serum (FBS) (1:2, Cultilab, Campinas, Brazil) or astrocyte medium (1:1, DMEM/F12 containing 10% FBS, 1% L-glutamine and 1% U/ml penicillin/streptomycin). Reactive astrocytes were transferred to 15ml tubes and centrifuged at 200g for 7 minutes. Then, the pellet was resuspended in conditioned medium enriched with growth factors (DMEM/F12 containing 2% supplement B27, 1% L-Glutamine, 1% Penicillin-streptomycin, 20ng/ml bFGF, 20ng/ml Heparin, and 10ng/ml EGF) and seeded at a cell density of 8×10^3^ cell/well, in a 24-well plate with a low attachment surface (poly-2-hydroxyethyl methacrylate, Sigma Aldrich, St. Louis, USA). After 24 hours, neurospheres were treated daily with LY450139 Semagacestat, Notch signaling inhibitor at five different working concentrations (treatment groups): 10 nM, 30 nM, 100 nM, 300 nM and 1 µM. We also included a control group for LY450139 Semagacestat’s vehicle (0.001% DMSO). Cells were incubated at 37°C in 95% CO_2_ incubator. After one week (eight days), neurospheres were imaged for analysis of area size.

### Immunohistochemistry/ Immunocytochemistry

For immunohistochemistry analysis, serial brain sections around the lesion core were rinsed several times with 1xPBS-0.1% Triton (0.1% PBST) and incubated in a blocking solution containing 5% normal goat serum NGS (5% NGS, 0.1% PBST) for 60 minutes, at room temperature. After the time, sections were incubated with selected primary antibodies previously included in blocking solution (5% NGS, 0.1% PBST) overnight, at 2–8°C. Primary antibodies: chicken anti-Glial fibrillary acidic protein, GFAP (AB5541, 1:1000, Millipore, Massachusetts, USA); rabbit anti-S100 calcium-binding protein β, S100β (AB868, 1:100, Abcam, Cambridge, USA); Rat anti-Galectin 3 protein, Gal3 (SC23938, 1:150, Santa Cruz, Texas, USA); mouse anti-Neurogenic locus notch homolog protein 1, Notch1 intra-cellular domain, NICD (AB128076, 1:500, Abcam, Cambridge, USA); rabbit anti-Gllioma-associated oncogene or Zinc finger protein, Gli1 (AB49314, 1:500, Abcam, Cambridge, USA); mouse anti-catenin beta-1 or β-catenin Subunit of the cadherin protein complex, β-catenin (05-665, 1:500, Millipore, Massachusetts, USA); rabbit anti-MKI67, KI67 (AB166667, 1:300, Abcam, Cambridge, USA); and rat anti-bromodeoxyuridine, BrdU clone RF06 (MCA6144, 1:500, Bio-Rad Laboratories, Hercules, USA). For BrdU labeling, the sections were incubated with 2N HCL solution for 30 minutes and neutralized with Borate Buffer (1 M, pH 8.5) before blocking step. The next day, sections were rinsed (0.1% PBST) and incubated with the corresponding secondary antibody in DAPI solution (1:10000, Sigma Aldrich, St. Louis, USA) for 60-90 minutes, at room temperature. Secondary antibodies: IgG goat anti chicken Alexa 488 (A11039, 1:500, Invitrogen, Massachusetts, USA); IgG goat anti mouse Alexa 488 (A11029, 1:500, Invitrogen, Massachusetts, USA); IgG goat anti rabbit Alexa 488 (A11008, 1:500, Invitrogen, Massachusetts, USA); IgG goat anti mouse Alexa 594 (A21125, 1:500, Invitrogen, Massachusetts, USA); IgG goat anti rat Alexa 594 (A11005, 1:500, Invitrogen, Massachusetts, USA); IgG goat anti chicken Alexa 647 (A-21449, 1:500, Invitrogen, Massachusetts, USA); and IgG goat anti rabbit Alexa 647 (A21245, 1:500, Invitrogen, Massachusetts, USA). Finally, the sections were rinsed and mounted onto slides with aqueous solution (Fluoromount G, Electron Microscopy Sciences, USA). Alternatively, for immunocytochemistry assays, cells were previously seeded on coverslips and submitted to *in vitro* astrocyte reactivation or were maintained under the same medium conditions as control. After 3 dpl, both control and reactive astrocytes were fixed in 4% PFA, rinsed several times with 1xPBS and permeabilized with 1xPBS-0.1% Triton (0.1% PBST) for 5 minutes and incubated in blocking solution containing 5% normal goat serum, NGS (5% NGS, 0.1% PBST) for 60 minutes, at room temperature. After the time, sections were incubated with selected primary antibodies previously included in blocking solution (5% NGS, 0.1% PBST) overnight, at 2-8°C. Primary antibodies: chicken anti-Glial fibrillary acidic protein, GFAP (AB5541, 1:1000, Millipore, Massachusetts, USA); and rabbit anti-MKI67, KI67 (AB166667, 1:300, Abcam, Cambridge, USA). The next day, sections were rinsed (0.1% PBST) and incubated with the corresponding secondary antibody in DAPI solution (1:10.000) for 60-90 minutes at room temperature. Secondary antibodies: IgG goat anti chicken Alexa 488 (A11039, 1:500, Invitrogen, Massachusetts, USA); and IgG goat anti rabbit Alexa 647 (A21245, 1:500, Invitrogen, Massachusetts, USA). Finally, the coverslips were rinsed and mounted onto slides with aqueous solution (Fluoromount G solution).

### Image acquisition

For both, *in vivo* and *in vitro* assays, image acquisition was performed using inverted microscope Olympus Life Science IX2 ILL100 (Tokyo, Japan) and at the confocal microscopy unit (INFAR, UNIFESP), using Leica TCS SP8 confocal microscope (Houston, USA) and Zeiss Observer Z.1. (Jena, Germany). Images were processed at maximum projection of the z-planes using ImageJ software (1.49v2, http://rsbweb.nih.gov/ij).

### Relative fluorescence intensity

We analyzed maximum projection images of at least three consecutive brain tissue sections around the lesion in 20x. For *in vitro* assays, we analyzed maximum projection images of ROIs in 20x of control and reactive astrocytes. We used ImageJ software. Channel images were transformed into 8 bits. Depending on the analysis we measured the correspondent mean gray value in the selected region of interest ROIs (3-5 fields/region, ipsilateral and contralateral). Values were normalized (maximum-minimum method) prior to statistical analysis.

### Overlapping pixels

We analyzed maximum projection images of at least three consecutive tissue sections around the lesion in 20x. We used ImageJ software. Channel images were trans-formed into 8 bits. After selecting ROIs around the lesion (3-5 fields/region, ipsilateral and contralateral), we established a minimum intensity value and threshold and used the histogram “count” tool to obtain the number of pixels by each channel. Next, we used the im-age calculator “and” tool to create a new image (mask) with the pixels shared in the two channels (GFAP/Gal3). Finally, we used the histogram “count” tool to obtain the number of pixels in the new image. Values were analyzed as a percentage of GFAP/Gal3 shared pixels of the total GFAP pixels by ROI.

### Signaling activation

We analyzed maximum projection images of at least three consecutive tissue sections around the lesion in 20x. We used ImageJ software. Channel images were trans-formed into 8 bits and inverted (black/white background). We used pseudocolor Look-Up table (LUT) images to differentiate intensity-labeling areas and a radial grid starting at the lesion site to determine the presence, distribution, and pattern of activation. A LUT is a predefined table of gray values with matching red, green, and blue values so that shadows of gray were displayed as colorized pixels. We applied a LUT scale (red-blue) macro to differentiate areas of high (red) and low (blue) immunolabeling. Additionally, we made a grade consisting of sequential concentric circles around the lesion using the tool “enlarge” and “Roi manager”. We set up 100µm between each new circle. Grades were used to determine the distribution (distance) of both high (red-core) and low (blue-periphery) signaling in LUT images.

### Nearest neighbor distance

Nearest neighbor distance (NND) is a method of description of the pattern of distribution of a population, adapted from Clark and Evans., (1954) work in ecology. The NND Ratio considers the mean distance between particles (objects, cells) and relative density. We analyzed maximum projection images of at least three consecutive tissue sections around the lesion in 20x. For *in vitro* assays, we analyzed maximum projection images of ROIs in 40x of control and reactive astrocytes. We used ImageJ software. GFAP channel images were transformed into 8 bits and were selected ROIs in the core and periphery around the lesion. For each ROI, we used “remove outliers” and “dilate” tools to create a new mask. Next were selected “centroid” and “area” in set measurements option and used the “analyze particles” tool (>20µm, mask outlines) to display the X and Y coordinates (centroid) of the particle. Finally, we used the NND plugin to get the nearest neighbor distances. Values were used to calculate the number of particles analyzed, density p (particles ⁄ total area ROI), mean area of the particles, mean distance, or mean expected distance RA and relative density RE (1⁄2√p). NND Ratio corresponded to RA ⁄ RE. Ratios close to zero describe a distribution of total aggregation, while ratios close to 2.15 describe a random distribution(https://icme.hpc.msstate.edu/mediawiki/index.php/Nearest_Neighbor_Distances_Calculation_with_ImageJ.html).

### Cell counting

We analyzed maximum projection images of at least three consecutive tissue sections around the lesion in 20x and 40x. For *in vitro* assays, we analyzed used maximum projection images of ROIs in 40x of control and reactive astrocytes. In the case of 20x were selected ROIs (3-5 fields/region, ipsilateral, contralateral). We used ImageJ software. For both 20x ROIs and 40x images, we used the “count cells” tool to quantify the total number of cells (Mature and & newly proliferating cells, DAPI), the number of newly proliferated cells Ki67, the number of GFAP reactive astrocytes, and the number of GFAP/Ki67 newly proliferated astrocytes. Ki67/DAPI cells corresponded to (+Ki67 cells) ⁄ (DAPI cells). Similarly, GFAP/Ki67 cells corresponded to (+GFAP+Ki67 cells) ⁄ (+GFAP cells). Results were presented as relative and absolute values.

### Connected component

Connected component analysis is a method to find and create “subgraphs” or connected components (groups of particles) based on the separation distance among particles (cells) in each assembling configuration. We analyzed maximum projection images of at least three consecutive tissue sections around the lesion in 40x. For *in vitro* assays we analyzed maximum projection images of ROIs in 40x of control and reactive astrocytes. We used ImageJ software. Ki67 channel images were transformed into 8 bits. For each image, we used “make binary” tool to create a new mask. Next were selected “area” and “centroid” in set measurements options and used the analyze particles tool (>10µm, record starts, mask outlines) to display the X and Y coordinates (centroid) and the XS and YS (edges) of the particles. Finally, we used the Graph plugin with 10µm between edges as assembling configuration. In the case of *in vitro* studies, we analyzed particles >20μm. Graph plugin was set at 40µm between edges. We obtained an adjacency list showing the number of connected components and descriptors. Values were used to quantify the number of connected components and the number of particles by components. We divided the list in-to two groups: a) a first, single group of up to two Ki67+ newly proliferated cells connected and, b) a second cluster group, with more than three Ki67+ newly proliferated cells connected. Results were presented as relative values (https://imagej.nih.gov/ij/plugins/graph/index.html).

### Neurosphere area size

We analyzed at least ten images per group in 20x. We used ImageJ software. Neuro-sphere area size was calculated using “area” tool. Values were used to compare the effect of Notch signaling inhibition treatment between groups.

### RNA extraction and cDNA synthesis

For each group, were used 5×10^5^ cells/sample approximately. Total RNA was extracted according to the manufactureŕs recommendation using PureLink RNA Micro Kit (12183-016, Invitrogen, Life Technologies, Massachusetts, USA). Samples were incubated in 1% 2-mercaptoethanol lysis solution and washed with 70% ethanol. Next, samples were transferred into columns, centrifuged, and were sequentially washed in buffer solutions I and II. Finally, samples were incubated in DNAse solution and resuspended in free-RNAse water. For each sample, total RNA was quantified by spectrophotometric assay using a NanoVue Plus spectrophotometer (GE Healthcare, Buckinghamshire, UK). RNA was reverse transcribed using High-Capacity cDNA Reverse Transcription Kit (4368814, Ap-plied Biosystems, Life Technologies, Massachusetts, USA). Briefly, 1-2 μg of total RNA was incubated in 2μL 10x RT Buffer, 2 μL 10x RT Random Primers, 0,8μL 25x dNTP Mix (100 mM), 1 μL Reverse Transcriptase 50 U/μL and 1 μL RNase Inhibitor using an Eppendorf Mastercycler Personal thermocycler system (Applied Biosystems, Foster City, USA). Thermal cycling conditions were 25°C, 10 minutes, 37°C, 120 minutes, and 85°C, 5 minutes.

### Quantitative RT-PCR and gene expression analysis

For Quantitative Real-Time PCR (qPCR), we used a mix of 4uL of cDNA (12.5 ng) and 5μL of Fast SYBR Green Master Mix (4385610, Applied Biosystems, Massachusetts, USA), 0.25 μL (250nM) of forward and reverse primers and 0.5uL free-RNAse water in an Ap-plied Biosystems 7500 Real-Time PCR System (Applied Biosystems, California, USA). Genes and primer sequences (forward, FW, reverse, RV): GAPDH, FW AG-GTCGGTGTGAACGGATTTG, RV TGTAGACCATGTAGTTGAGGTCA; GFAP, FW GTTAAGCTAGCCCTGGACATC, RV GATCTGGAGGTTGGAGAAAGTC; S100β, FW AGAGGGTGACAAGCACAAG, RV CCACTTCCTGCTCCTTGATT; NESTIN, FW CTCAACCCTCACCACTCTATTT, RV CTGTGGCTGCTTCTTTCTTTAC; KI67, FW ATCATTGACCGCTCCTTTAGGT, RV GCTCGCCTTGATGGTTCCT; OLIG2, FW TCCCCAGAACCCGATGATCTT, RV CGTGGACGAGGACACAGTC; MASH1, FW CTT-GAACTCTATGGCGGGTT, RV TAAAG TCCAGCAGCTCTTGTT; HES1, FW CTATCATGGAGAAGAGGCGAAG, RV CCGGGAGCTATCTTTCTTAAGTG; HEY1 FW CCGACGAGACCGAATCAATAAC, RV TCAGGTGATCCACAGTCATCTG; CCND1, FW GATGAAGGAGACCATTCCCTTG, RV TCACCAGAAGCAGTTCCATTT. Thermal cycling conditions were 95°C for 20 seconds, 40 × 95°C for 3 seconds, and 58°C for 30 seconds. The dissociation curve was performed at 95°C for 1 minute, 60°C for 30 seconds, and 95°C for 30 minutes. We considered three to five biological replicates for each group and three technical replicates for each gene, and GAPDH gene in each plate. Gene expression was normalized to GAPDH expression, and the ΔCT method (Livak et Schmittgen, 2001) was used for relative quantification analysis.

### Statistical Analysis

For *in vivo* and *in vitro* studies, were considered both absolute and relative values. For *in vitro* gene expression analysis, the values were transformed using the ΔCT method (Livak et Schmittgen, 2001). Additionally, values in the fold change analysis corresponded to Log_2 (x+1) transformation of gene expression (ΔCT) values. Depending on the case, the graphs show mean, standard deviation (SD) and standard error (SEM). Statistical significance was evaluated using one-tailed un-paired Student’s t test for 2-group comparison, and Analysis of Variance (ANOVA) and Bonferroni pos-hoc test for multiple comparison. Kruskal-Wallis test was used to evaluate the effect of Notch inhibitor doses over neurosphere area size. Values with a confidence interval of 95% (p < 0.05) were considered statistically significant. Statistical analysis of the data and the graphical representations were performed using GraphPad Prism (v 5.0, San Diego, USA) software and Excel (v. 2018, https://www.microsoft.com/pt-br/microsoft-365/excel) software.

